# Nonlinear post-selection inference for genome-wide association studies

**DOI:** 10.1101/2020.09.30.320515

**Authors:** Lotfi Slim, Clément Chatelain, Chloé-Agathe Azencott

## Abstract

To address the lack of statistical power and interpretability of genome-wide association studies (GWAS), gene-level analyses combine the p-values of individual single nucleotide polymorphisms (SNPs) into gene statistics. However, using all SNPs mapped to a gene, including those with low association scores, can mask the association signal of a gene.

We therefore propose a new two-step strategy, consisting in first selecting the SNPs most associated with the phenotype within a given gene, before testing their joint effect on the phenotype. The recently proposed kernelPSI framework for kernel-based post-selection inference makes it possible to model non-linear relationships between features, as well as to obtain valid p-values that account for the selection step.

In this paper, we show how we adapted kernelPSI to the setting of quantitative GWAS, using kernels to model epistatic interactions between neighboring SNPs, and post-selection inference to determine the joint effect of selected blocks of SNPs on a phenotype. We illustrate this tool on the study of two continuous phenotypes from the UKBiobank.

We show that kernelPSI can be successfully used to study GWAS data and detect genes associated with a phenotype through the signal carried by the most strongly associated regions of these genes. In particular, we show that kernelPSI enjoys more statistical power than other gene-based GWAS tools, such as SKAT or MAGMA.

kernelPSI is an effective tool to combine SNP-based and gene-based analyses of GWAS data, and can be used successfully to improve both statistical performance and interpretability of GWAS.

## 1. Introduction

Lack of statistical power is a major limitation of genome-wide association studies (GWAS).If we perform tests at the level of single nucleotide polymorphisms (SNPs), this lack of statistical power stems from from small effect sizes and linkage disequilibrium, among others. By modeling the association signal over an entire region, gene-level analyses can address this limitation. Being functional entities, genes have the potential to shed light on yet undiscovered biological and functional mechanisms. Indeed, several tools have been proposed in recent years to aggregate SNP-level information into gene statistics.^1–3^ However, using all SNPs mapped to a gene, including those with low association scores, can mask the association signal of a gene. We therefore propose a new two-step strategy, consisting in (1) restricting ourselves to the SNPs most associated with the phenotype within a given gene, and (2) testing their joint effect on the phenotype.

Because of the data scarcity in GWAS, we would like to use the same data for these two steps. However, without further precaution, this amounts to data snooping, and may lead to overestimating the effect of the gene on the phenotype. Post-selection inference (PSI)^4^ is a framework specifically developed to address this issue. The main idea is to correct for selection bias by conditioning the models used in the second step on the selection event from the first step.

We have previously proposed a generic framework for nonlinear post-selection inference called kernelPSI.^5^ kernelPSI uses quadratic kernel association scores (QKAS) to perform the selection step. They are quadratic forms of the response vector, and can measure nonlinear association between a group of features and the response. In the context of GWAS, QKAS can be used to model epistatic interactions between neighboring SNPs, and the kernelPSI framework is therefore well-suited to performing our two-step strategy.

In this paper, we show how to use kernelPSI in the GWAS setting. The first step consists in selecting, for each gene or region of interest, a number of putative loci, or blocks of loci, according to an appropriate QKAS. The second step tests for their aggregated effect on the phenotype. The selection bias introduced by the first step is accounted for by sampling constrained replicates of the response vector. The statistics of the response are compared to those of the replicates to obtain valid *p*-values.

The extension of kernelPSI to GWAS required several modifications. First of all, we generalized kernelPSI to any non-normally distributed continuous outcome, enabling us to use it on any continuous phenotype. Second, we used contiguous hierarchical clustering^6^ to separate each gene in blocks of strongly correlated SNPs. Third, we implemented the identical-by-state kernel^1^ to define the similarity between individuals based on the number of individual alleles they share in such a block of SNPs. Finally, we substantially improved the scalability of the kernelPSI code. Most importantly, we developed a GPU-version of the constrained sampling algorithm to speed-up linear algebra operations. The rest of the code was also accelerated thanks to a more efficient C++ backend. In particular, we implemented a rapid estimator of the HSIC criterion^7^ based on quadratic-time rank-1 matrix multiplications. HSIC is an example of quadratic kernel association score.^5^

To illustrate the use of kernelPSI on GWAS data, we present a study of BMI and its fluctuations (ΔBMI) in the UK BioBank.^8^ The UK BioBank is one of the largest available sources of data for the investigation of the contribution of genetic predisposition to a variety of physiological and disease phenotypes. We study both BMI and ΔBMI because of the suspicion that different genetic mechanisms might be governing the two phenotypes.^9^ Our study yielded a list of causal *loci* within genes playing a putative role in BMI and ΔBMI. In addition, this use case also shows that kernelPSI enjoys a better statistical performance than other gene-level association tools, with the unique benefit of pinpointing the causal *loci* within these genes. We propose an eponymous R package that implements the full pipeline of kernelPSI. The CPU-only version is directly available from CRAN. The enhanced GPU-version can be downloaded from the development branch of the GitHub repository https://github.com/EpiSlim/kernelPSI.git.

## 2. Methods

### 2.1. kernelPSI for GWAS

We describe here how to use the kernelPSI framework^5^ to test for association between genetic variants in a region and a continuous trait.

#### 2.1.1. Notations

We assume *n* samples are sequenced in a region containing *p* SNPs. For each sample *i* ∈ ⟦1, *n*⟧, *y_i_* ∈ ℝ represents the phenotype and *x_i_* the genotypes for the *p* SNPs. We denote by *y* ∈ ℝ^*n*^ the vector of phenotypes. Assuming a dosage encoding (or an additive genetic model), *x_ij_* ∈ {0, 1, 2} represents the number of copies of the minor allele at locus *j* for sample *i*. Other genetic models and encodings can also be considered; dominant and recessive models, for example would results in *x_ij_* ∈ {0, 1}.

We further consider a partition of the region in a set of *S* contiguous SNP clusters 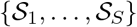. Such clusters correspond to LD blocks and their construction is detailed in Section 2.3. For any *i* ∈ ⟦1, *n*⟧ and *t* ∈ ⟦1, *S*⟧, we denote by ***x**_i,t_* the vector ***x**_i_* restricted to the SNPs in cluster 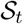, that is to say, the values of the SNPs of cluster 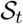 for sample *i*. Our goal is to first select the SNP clusters most associated with the phenotype, and second measure the association of the entire genomic region with the phenotype through its joint association with the selected clusters.

#### 2.1.2. Scoring clusters with kernels

We propose to use kernels to perform the selection step. More specifically, we use a positive semi-definite kernel function *k_d_* : {0, 1, 2}^*d*^ × {0, 1, 2}^*d*^ → ℝ (defined for any *d* ∈ ℕ∗) to define, for each cluster 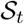, a kernel matrix *K_t_* ∈ ℝ*n*×*n* such that, for any *i, k* ∈ ⟦1, *n*⟧, 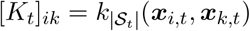. Intuitively, [*K_t_*]_*ik*_ is a measure of the similarity between samples *i* and *k*, based on the 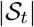 variants in the *t*-th cluster of the genomic region. One of the most commonly used kernel function in genomics is the weighted IBS kernel, which is described in more details in Section 2.4.

Having defined these kernel matrices, we select the kernels most associated with the phenotype thanks to a so-called *quadratic kernel association score,* that is to say, a scoring function of the form

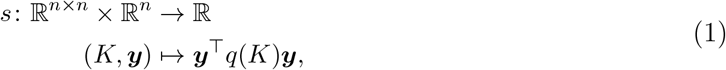

for a mapping *q* : ℝ^*n*×*n*^ → ℝ^*n*×*n*^. We can use such a score to quantify the association between a single cluster and the phenotype as *s*(*K_t_, **y***), but also to quantify the association between a set of clusters 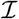 and the phenotype as 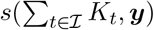.

Quadratic kernel association scores are quite generic, and encompass a number of propositions. For example, using the identity for *q* is equivalent to the Sequence Kernel Association Test (SKAT) score.^1^ In this paper, and because this choice was showed to lead to improved statistical power in a variety of settings,^5^ we use the unbiased Hilbert-Schmidt Independence Criterion (HSIC) estimator.^10^ HSIC has been in a variety of bioinformatics applications in the past,^11^ including the non-linear analysis of GWAS data.^12,13^ The unbiased HSIC estimator between two kernel matrices *K* and *L* is given by:

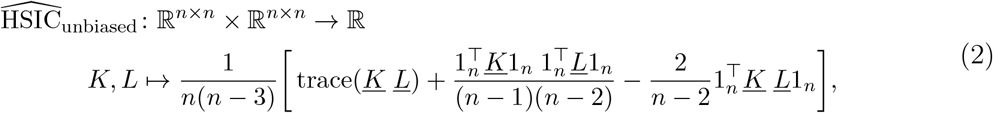

where 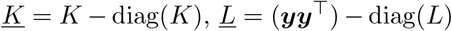, and 1_*n*_ is a *n*-dimensional vector of all ones. We simply define

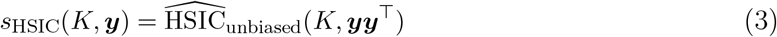

We refer the reader to our earlier work^5^ for the proof that this estimator is indeed a quadratic kernel association score.

#### 2.1.3. Adaptive forward selection strategy

Several strategies for cluster selection can then be used to perform selection based on a choice of quadratic kernel association score. In particular, one can adopt a forward- or backward-stepwise strategy, and the number of selected clusters can be fixed or adaptively determined. We opt here for an adaptive forward strategy, which consists in iteratively adding clusters to the (initially empty) set of selected clusters until it contains all clusters, and then choosing the number of selected clusters *S*∗ according to the maximum score attained throughout the procedure. The detailed algorithm can be found in Appendix A.1.

#### 2.1.4. Post-selection inference

We now want to use the optimal quadratic kernel association score *s*∗ to determine the association of the genomic region with the phenotype *y,* through its association with the selected clusters in 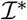. To obtain valid p-values, we need to account for the fact that we performed this selection step, and that the clusters were chosen for their strong score of association with *y*.

In other terms, denoting by 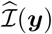 the set of indices of clusters selected by our procedure for a phenotype ***y***, we need to determine the distribution of 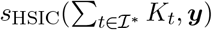 *conditionally* to the selection event 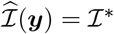.

We have shown^5^ that this selection event can be modeled as an intersection of a finite number *M* = (2*S* − 1) of quadratic constraints on ***y***. Unfortunately, this does not allow us to determine the exact distribution of *s*_HSIC_ conditionally to this selection event. However, the sampling procedure we proposed in^5^ makes it possible to efficiently sample replicates of the outcome ***y*** which statisfy those quadratic constraints, which we leverage to determine empirically the null distribution and obtain the desired p-values.

### 2.2. Outcome normalization

Our original proposal5 is limited to normally-distributed outcomes. To expand kernelPSI to other continuous outcomes, it is sufficient to transform any continuous outcome *y* into a vector of independent normally-distributed variables. Several outcome normalization approaches have been proposed, among which we found the van der Waerden quantile transformation to be the most consistent approach across phenotype distributions. We provide more details in Appendix A.3.

### 2.3. Contiguous hierarchical clustering for genomic regions

We could apply the strategy outlined in Section 2.1 without any clustering step, using *p* clusters of 1 SNP, so as to determine the association of a genomic region with the phenotype through the association of individual SNPs.

However, nearby SNPs are often strongly correlated, a phenomenon refered to as linkage disequilibrium (LD). Using clusters of correlated variants rather than individual SNPs has the double advantage of reducing the number of clusters to choose from, while simultaneously modeling the combined effects of the SNPs within a cluster on the phenotype. It also reduces the dependence between individual tests.

To define these clusters, we use the R package BALD^6^ which implements adjacent hierarchical clustering (AHC) in conjunction with the gap statistic^14^ to determine the optimal number of clusters *S*. This approach is illustrated on Supplementary Figure 1.

### 2.4. The IBS-kernels and nonlinear SNP selection

We must now choose a kernel function *k_d_* that we will use to construct the kernel matrices *K_t_* corresponding to each of the clusters 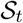, and interpreted as a similarity between two samples based on the SNPs of cluster 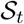. For SNP data, the identitcal-by-state (IBS) kernels proposed by Wu et al.,^15^ which use the number of alleles that are identical between individuals, is a popular choice. We give more details about these kernels in Appendix A.4.

### 2.5. Efficient nonlinear post-selection inference for high-dimensional data

In this section, we detail a number of modifications we included in order to improve the scalability of kernelPSI to the large sample sizes.

#### 2.5.1. Rapid HSIC estimation

We score the association between a cluster and the phenotype using the HSIC unbiased estimator (see Equation (3)). We recall here Equation (2):

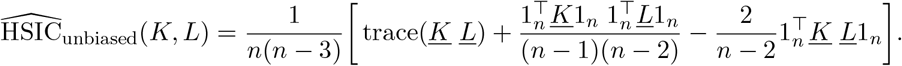

The computation of 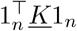 and 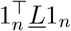 can be performed in quadratic time 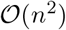. However, computing trace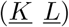 and 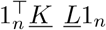 requires the matrix-matrix multiplication of 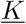 and 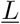, resulting in a 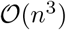 complexity. To avoid that, we decompose trace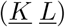 as 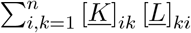, which results in a better 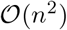 complexity. The same complexity can be achieved for the quadratic form 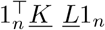 by starting with the matrix-vector multiplication of either 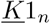 or 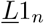. Overall, we achieve a 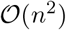 complexity, for which 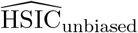 can be computed on a single CPU for thousands of samples in relatively little time. As an illustration, we performed 100 repetitive evaluations of 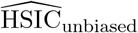 for two matrices of size 5, 000 × 5, 000. On a 2.7 GHz intel core i5 processor, the average running time was 1.08s.

#### 2.5.2. Accelerated replicates sampling

The gains achieved by speeding up the estimation of 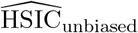 turned out to be insufficient because of the heavy computational workload involved in sampling replicates to compute p-values. Our sampling algorithm^5^ is partly a rejection sampling algorithm. At every iteration, we verify that the candidate replicate satisfies *M* quadratic constraints ***y***^T^*Q_m_**y*** + *b_m_* ≥ 0 for *m* = 1*, …, M*. In Appendix A.6, we show in simulations that the *p*-values obtained through our sampling algorithm are statistically valid. The kernelPSI statistics are not overpowered, since the *p*-values are uniformly distributed under the null hypothesis (absence of effects). Furthermore, kernelPSI has statistical power under the alternative hypothesis.

When *M* is large, we observed a significant slow-down due to the overhead between successive evaluations of the constraints. A single combined evaluation eliminates this overhead. We achieve this by encoding all computations in a matrix form, as illustrated on Supplementary Figure 3, and by using the ViennaCL library^16^ to accelerate these computations on GPUs.

A major drawback in hybrid CPU-GPU calculations is the transfer time between the main memory and the GPU memory. With most Nvidia GPUs, the theoretical bandwidth limit is 8 Gb/s. To reduce transfer times of the matrices 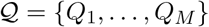, we transfer them to GPU memory once and for all before sampling. However, because of memory size limitations, this imposes an upper limit on the number *M* of matrices in 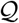, and consequently on the number of LD clusters as *S* = *M/*2 + 1.

Finally, we give a rough estimation of the complexity of our sampling algorithm. If we denote by *N*_replicates_ the number of replicates, the overall complexity can be approximated as 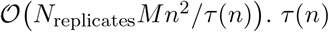 is a decreasing function of *n* which corresponds to the probability of sampling a replicate in the acceptance region. The average number of iterations to obtain a valid replicate is then 1/*τ* (*n*) (mean of a geometric distribution). We are currently unable to propose a closed form for *τ* (*n*).

### 2.6. Data and experiments

The study of physiological phenotypes in GWAS has so far focused on basic anthropometric measures such as height, weight, and BMI. Their longitudinal fluctuations received little attention, mainly because of the lack of such data. To the best of our knowledge, the fluctuations of BMI in adults have been the subject of very few GWA studies.^17–19^ Some studies^9^ suggest that BMI and its variation might be influenced by distinct sets of SNPs. In addition, rare variants impacting weight loss through gene-diet interaction are referenced in the literature.^20^ Recent biobanks are finally making such longitudinal data available, among which the UK BioBank,^8^ which provides extensive phenotypic and health-related information for over 500, 000 British participants. We apply kernelPSI on the UK BioBank dataset to separately study BMI and variations of BMI (ΔBMI).

#### 2.6.1. Quality control

Our preprocessing pipeline for the UK BioBank dataset is close to the pipeline of the Neale lab^a^. The detailed quality control procedure we conducted for samples can be found in Appendix A.7. The application of this pipeline yielded *n* = 266, 679 final samples. In contrast, the Neale lab obtained 337, 000 samples by implementing less stringent thresholds. The QC pipeline resulted in 577, 811 SNPs.

#### 2.6.2. Phenotypes

ΔBMI is not directly available. We computed it from the participants who attended both the initial assessment visit and the first repeat assessment visit. In total, we obtained 11, 992 samples for ΔBMI. More precisely, we define ΔBMI as the average yearly variation (BMI_repeat_ − BMI_initial_)/Δ*t*_years_. Indeed, the time span between the visits is not the same for all participants.

As for BMI, we use the measurements of the initial visit. To avoid issues that can arise when jointly analysing results stemming from datasets containing different samples, we restrict ourselves to studying the samples for which both phenotypes are available.

As explained in the Section 2.2, we apply the Van der Waerden transformation to the phentoypes. We show on Supplementary Figure 2 a visual comparison of the empirical cumulative distribution functions (cdfs) of BMI and ΔBMI to the cdf of a standard normal distribution.

#### 2.6.3. Gene selection

To reduce computational time and resource requirements, as well as facilitate results validation, we restricted our study to the genes already associated with BMI in the GWAS catalog.^21^ The scope of the narrower study is then gene prioritization: the dual SNP-gene perspective of kernelPSI allows it to assess whether SNP-level associations translate into a gene-level association. This is particularly interesting given the large number of genes associated with BMI (1811 genes). For each gene, the sampling of 50, 000 replicates took on average 6 minutes on a single K80 Nvidia GPU. This makes kernelPSI scalable to genome-wide analyses.

To define genic boundaries, we used the biomaRt tool,^22^ which provided a genomic interval for 1774 genes. Moreover, the intervals were converted from the GRCh38 coordinate system to the GRCh37 one, since the SNP positions in the UK BioBank are given in the GRCh37 system. We point out that the conversion can result in several noncontiguous intervals.^23,24^

An immediate use of the resulting intervals led to a number of genes without any SNPs within. As a result, we added a downstream/upstream 50kb buffer to cover more SNPs. The same buffer size was also opted for by several other authors.^25,26^

#### 2.6.4. SNP clusters

Despite the extended 50kb buffer, several genes still contained a handful of SNPs. 1215 genes contained at most 3 SNPs. In particular, if only one SNP is mapped to a given gene, the kernel selection step becomes irrelevant. Nonetheless, we still perform hypothesis testing by directly using the HSIC statistic in Eq. (2) to measure the association between the gene and the phenotype. If 2 or 3 SNPs are mapped to a gene, we associate a distinct cluster/kernel to each one of them. This allows for a more accurate SNP selection.

For all other genes (i.e. those mapped to more than 4 SNPs), we applied AHC, as explained in Section 2.3. The optimal number of clusters *S* is determined by the gap statistic. In the adaptive kernel selection strategy we use here, this leads to *i_M_* = 2 (*S* − 1) constraints. To avoid the issues encountered for large values of *i_M_* (see Section2.5.2), we set the maximum number of clusters to 5. This leads to a ∼ 9.2Gb maximum GPU memory occupancy for 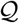.

## 3. Results

### 3.1. Kernel selection

Validating statistical tools in GWAS is always made difficult by the lack of a ground truth. In our study, validation is made easier by the fact that we only tested genes containing SNPs previously associated to BMI in the GWAS catalog. We can therefore evaluate kernelPSI based on its ability to recover GWAS catalog SNP, which we measure using the distance between those SNPs and the clusters selected by kernelPSI. Figure 1 shows the distribution histogram of these distances. It is heavily skewed toward small distances, meaning that the clusters selected by kernelPSI are often located near the GWAS catalog SNPs. This confirms the capacity of kernelPSI to retrieve relevant genomic regions. Moreover, the selected clusters also surround GWAS catalog SNPs. For BMI and ΔBMI, the selected clusters respectively included at least one GWAS catalog SNP in 62.5% and 40.6% of genes.

**Fig. 1.**
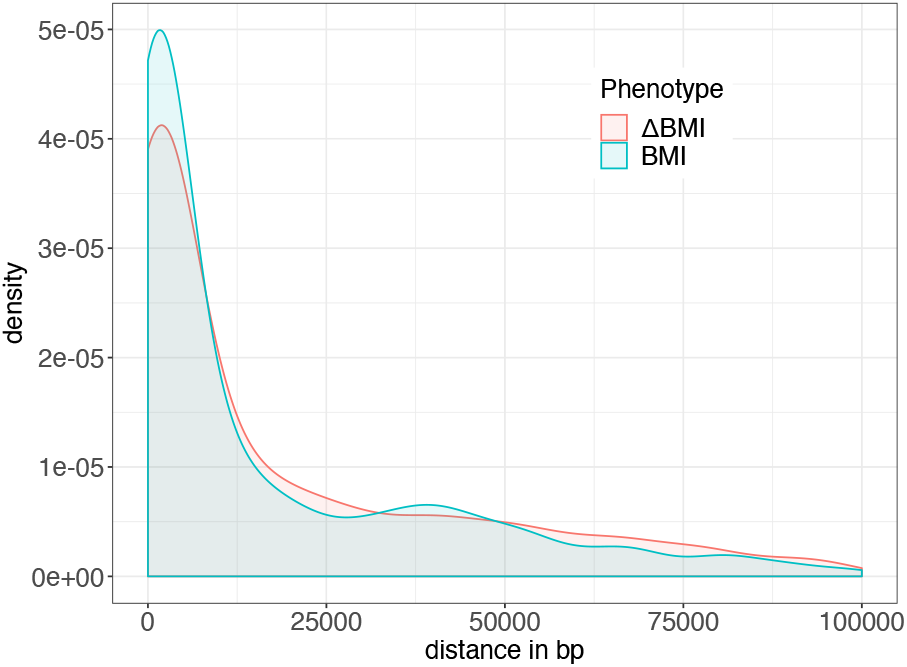
Distance between the SNPs of the GWAS Catalog and their closest neighbor among the SNPs in the clusters selected by kernelPSI.

**Fig. 2.**
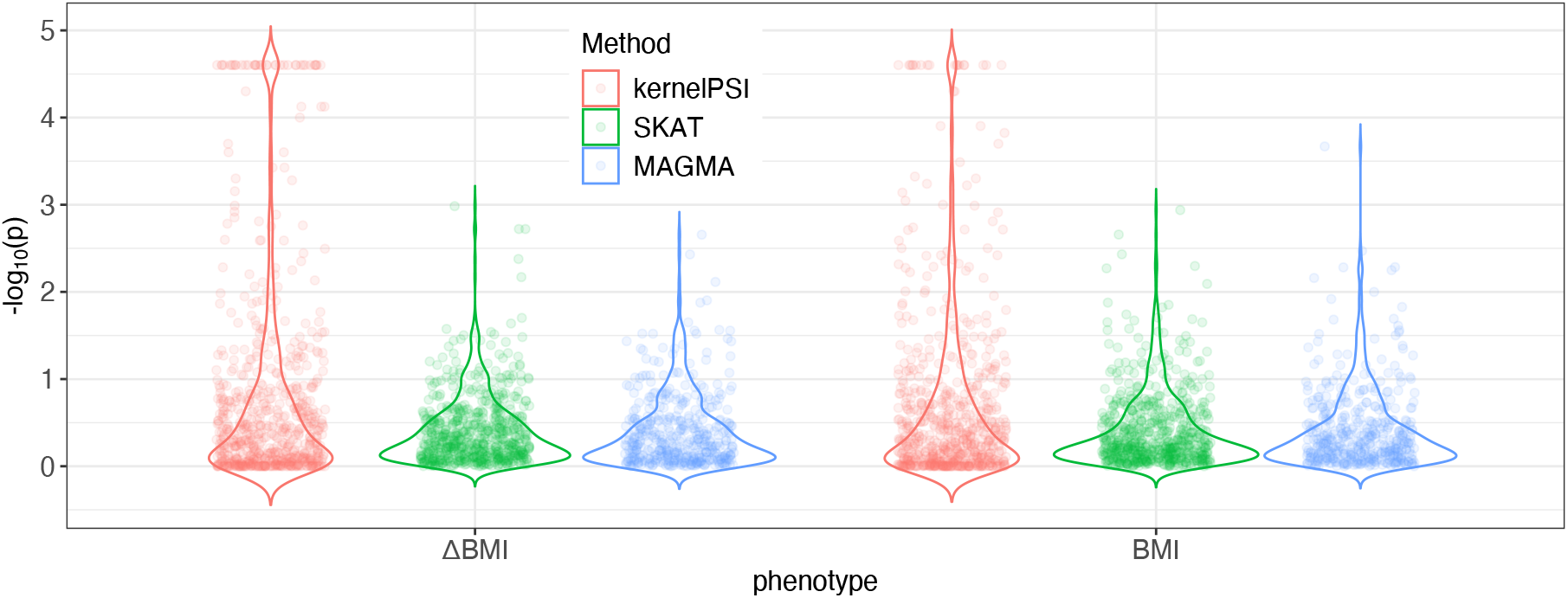
Violin plot comparing the *p*-values of kernelPSI for BMI and ΔBMI to two benchmarks.

These results would be irrelevant if kernelPSI turned out to be selecting all clusters. Indeed, the clusters would always contain GWAS catalog SNPs. However, kernelPSI was conservative for a large majority of genes, overwhelmingly selecting fewer clusters than the total number of clusters *S* (see Table 1). For BMI, kernelPSI selected one cluster in 75, 9% of the genes for which *S* = 3 and at most 2 clusters in 73, 6% of the genes for which *S* = 5. Similar results were obtained for ΔBMI.

**Table 1.** Distribution of the number of selected clusters *S*^∗^ depending on the total number of clusters *S* and the phenotype.

|  | BMI |  |  |  |  | ΔBMI |  |  |  |  |
| --- | --- | --- | --- | --- | --- | --- | --- | --- | --- | --- |
| | $S = 1$ | $S = 2$ | $S = 3$ | $S = 4$ | $S = 5$ | $S = 1$ | $S = 2$ | $S = 3$ | $S = 4$ | $S = 5$ |
| $S^* = 1$ | 100.0% | | | | | 100.0% | | | | |
| $S^* = 2$ | 82.9% | 17.1% | | | | 92.7% | 7.3% | | | |
| $S^* = 3$ | 75.9% | 18.5% | 5.6% | | | 70.4% | 20.4% | 9.3% | | |
| $S^* = 4$ | 55.9% | 30.9% | 10.3% | 2.9% | | 50.0% | 27.9% | 19.1% | 2.9% | |
| $S^* = 5$ | 43.9% | 29.7% | 17.6% | 7.2% | 1.5% | 40.3% | 28.4% | 21.4% | 7.6% | 2.3% |

Overall, the conservative kernel selection combined with the proximity of the selected kernels to the GWAS catalog SNPs demonstrate the selective abilities of kernelPSI.

### 3.2. Hypothesis testing

For association testing, we benchmarked kernelPSI against two state-of-the-art gene-level baselines. The first one is SKAT,^1^ and can be described as a non-selective variant of kernelPSI. Furthermore, it is a quadratic kernel association score which can be incorporated into the framework of kernelPSI. The SKAT score is a variance-component score^27^ given by *s*_SKAT_(*K, Y*) = *Y*^T^*KY*, for a centered phenotype *Y*. The second baseline is MAGMA,^2^ which computes an F-test in which the null hypothesis corresponds to absence of effects of all genotype PCs.

To compute the empirical *p*-values in kernelPSI, we sampled 40, 000 replicates in addition to 10, 000 burn-in replicates. The comparison of the distributions of the resulting *p*-values to those of SKAT and MAGMA shows that kernelPSI clearly enjoys more statistical power than the two baselines for both phenotypes (Figure 3.2). The *p*-values were altogether significantly lower. Thanks to the large number of replicates, we attribute this performance, not to the lack of accuracy of the empirical *p*-values, but to the discarding of non-causal clusters in the selection stage.

Our study was motivated by the hypothesis that BMI and ΔBMI are driven by different biological mechanisms. The low rank correlations of the *p*-values between the two phenotypes (see Table 2) lend further credence to this hypothesis. Interestingly, we observed a similar range of values for kernelPSI and the two benchmarks SKAT and MAGMA. For all metrics and methods, the rank correlations are lower than 0.1.

**Table 2.**
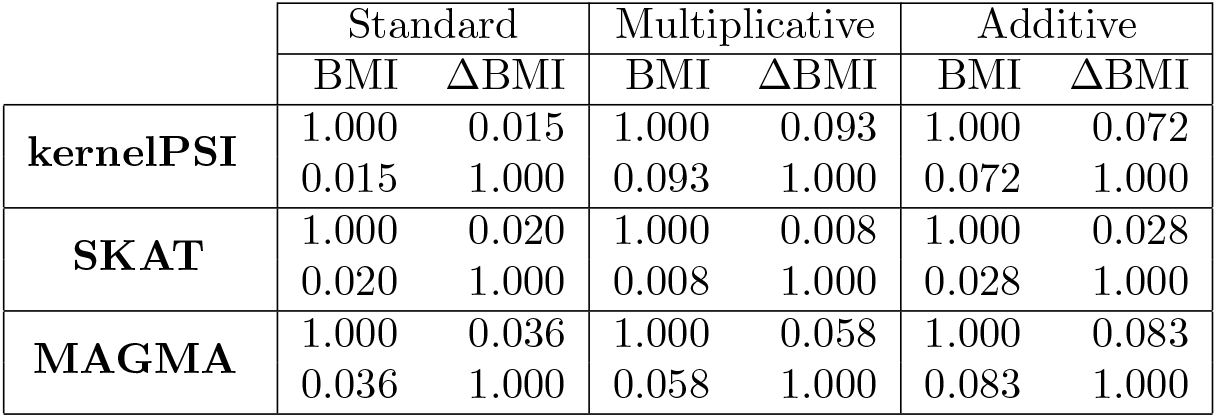
Concordance between the *p*-values for BMI and ΔBMI by method, according to three Kendall rank correlation measures (standard, multiplicative, additive)

|  | Standard |  | Multiplicative |  | Additive |  |
| --- | --- | --- | --- | --- | --- | --- |
| | BMI | ΔBMI | BMI | ΔBMI | BMI | ΔBMI |
| **kernelPSI** | 1.000 | 0.015 | 1.000 | 0.093 | 1.000 | 0.072 |
|  | 0.015 | 1.000 | 0.093 | 1.000 | 0.072 | 1.000 |
| **SKAT** | 1.000 | 0.020 | 1.000 | 0.008 | 1.000 | 0.028 |
|  | 0.020 | 1.000 | 0.008 | 1.000 | 0.028 | 1.000 |
| **MAGMA** | 1.000 | 0.036 | 1.000 | 0.058 | 1.000 | 0.083 |
|  | 0.036 | 1.000 | 0.058 | 1.000 | 0.083 | 1.000 |

Despite the low rank correlations between BMI and ΔBMI, we obtained 7 common significant genes (*CKB, EIF2S2, KSR2, MIR100HG, NRXN3, PDILT* and *RAB27B*) out of 64 significant genes for ΔBMI and 40 for BMI. The latter were determined after the application of the Benjamini-Hochberg procedure with an FDR threshold of 0.05. The existence of a number of separate mechanisms does not preclude the existence of common ones simultaneously regulating BMI and ΔBMI.

## 4. Conclusion

Several tools have already been proposed to address the lack of statistical power of SNP-based GWAS by performing analyses at the gene level. However, aggregating into gene p-values information from all SNPs mapped to a gene, including those with low association scores, can weaken the association signal. We therefore proposed a two-step strategy, consisting in (1) restricting ourselves to the SNPs most associated with the phenotype within a given gene, and (2) testing their joint effect on the phenotype.

Recent advances in the field of post-selection inference make it possible to perform these two steps on the same data, and to obtain valid p-values without overestimating the joint effect of the selected SNPs on the phenotype. In addition, one can model nonlinear effects and interactions among SNPs within this framework, thanks to the use of kernels.

In this paper, we have showed how to use the generic post selection inference framework kernelPSI in the GWAS setting. To this end, we included several modifications, generalizing kernelPSI to non-normally distributed continuous outcomes, using adjacent hierarchical clustering to separate each gene in blocks of strongly correlated SNPs, implementing the identical-by-state kernel to define similarities between individuals, and substantially improving the scalability of the code.

Using data from the UKBiobank, we have illustrated how these modifications make it possible to analyze two continuous phenotypes, BMI and its variation. We have shown on this case study that kernelPSI enjoys more statistical power than either SKAT or MAGMA, and successfully identifies genes associated with a phenotype through the signal carried by the most strongly associated regions of these genes.

The broad GWAS community can benefit from tools like kernelPSI which combine statistical performance with interpretability. In the future, we hope that developments in the post-selection inference field will allow us to develop exact variants of kernelPSI, which forego the sampling step to directly determine the associated *p*-value. This could further reduce computational times dramatically.

Another direction of major interest for statistical genetics would be the application of kernelPSI to other SNP sets, such as cis-regulatory regions, or entire pathways. Indeed, it should be possible to use kernelPSI to determine pathways significantly associated with a phenotype, through the association signal carried by individual genes or even LD-blocks of each pathway.

## Supplementary Materials

Appendices are available at https://doi.org/10.1101/2020.09.30.320515

## Acknowledgments

This work used the UK Biobank Resource (Application Number 48319), and was supported by Agence Nationale de la Recherche (ANR-18-CE45-0021-01 and ANR19-P3IA-0001).

## A Supplementary Materials

## A.1 Adaptive forward selection algorithm

Our adaptive forward selection algorithm is as follows:

- **Initialisation**:

– A set 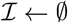 of indices of selected clusters
– A set 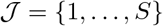 of indices of non-selected clusters
– An optimal set of indices of selected clusters 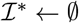 and an optimal score *s*^∗^ ← 0
- **for** *i* = 1 to *S* :

– 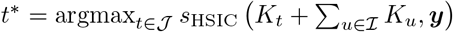
– 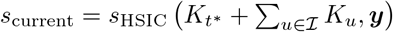
– 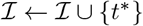
– 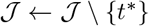
– **if** *s*_current_ > *s*^∗^:

* *s*∗ ← *s*_current_
* 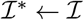
- **return** 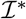

## A.2 Adjacency-constrained hierarchical clustering

Figure 1 illustrates the adjacency-constrained hierarchical clustering procedure to determine clusters of SNPs in LD.

## A.3 Outcome normalization

The van der Waerden outcome normalization is given by

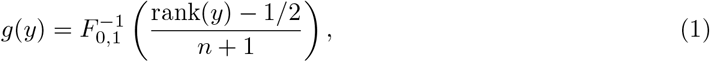

where *y* ∈ ℝ, rank(*y*) is the ranking of *y* in a descending order with respect to *y*_1_*, …, y_n_*, and *F*_0,1_ is the cumulative distribution function of the standard normal distribution.

The accuracy of the transformation in Equation (1) depends on the accuracy of the estimation of the regularized quantile (rank(*y*) - 1*/*2)*/*(*n* + 1), and thus on the number of participants *n*. Thankfully, many recent GWAS, in particular for physiological measurements, boast a large number of participants.

**Figure S1:**
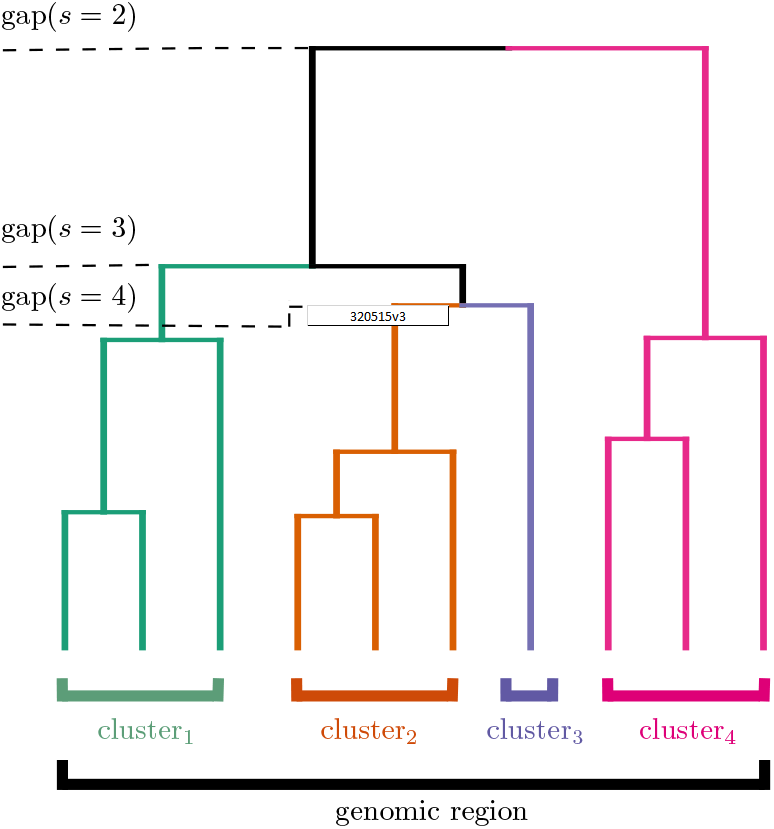
Clustering methodology: adjacent hierarchical clustering coupled with the gap statistic to determine the appropriate number of clusters.

Other outcome normalization methods have been proposed, such as the Lambert *W × F* [1], or Box-Cox [2] and Yeo-Johnson [3] transformations. In practice, we found the Van der Waerden transformation in Equation (1) to be the most consistent approach across different types of outcome distributions. All the above transformations are implemented in the R package bestNormalize [4].

Figure 2 illustrates the quality of the van der Waerden quantile normalization of the phenotypes. We notice a complete overlap between the cdfs. We attribute this good performance to the total number of samples and the low number of ties (11, 933 unique values).

**Figure S2:**
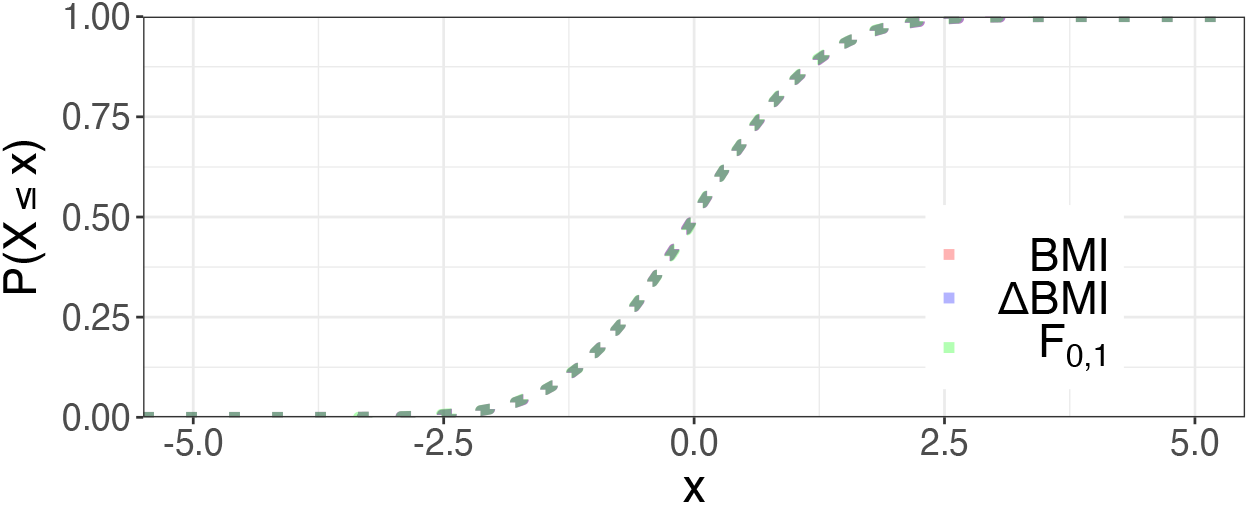
Comparison of the empirical cdfs of BMI and ΔBMI, respectively, to the cdf of a standard normal distribution.

## A.4 Identical-By-State Kernels

The most simple choice of kernel function would be the linear kernel *k_d_* : (***x**, **x***′) ↦ ⟨***x**, **x***′⟩. However, this does not take into account minor allele frequencies (MAFs) nor epistatic interactions between SNPs. To address this limitation, Wu et al. [5] proposed identical-by-state (IBS) kernels, which use the number of identical alleles between two individuals :

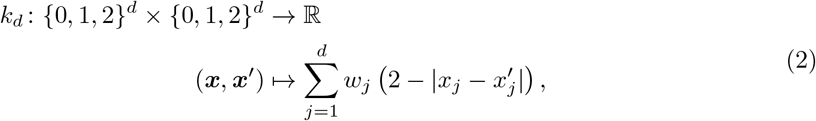

where the weight *w_j_* associated with SNP_*j*_ is a function of its MAF *m_j_* :

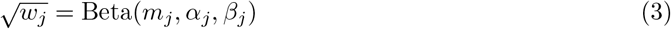

with Beta the density function of the Beta distribution. The idea of this parametrization is to given more importance to rare variants. The values of *α_j_* and *β_j_* are chosen according to the scope of the GWAS study. For common variants, Ionita-Laza et al. [6] recommend setting *α_j_* = *β_j_* = 0.5. To take an example, this assigns a weight of *w_j_* = 0.4 to a SNP of MAF 0.5 and a weight of *w_j_* = 10.2 to a SNP of MAF 0.01.

**Figure S3:**
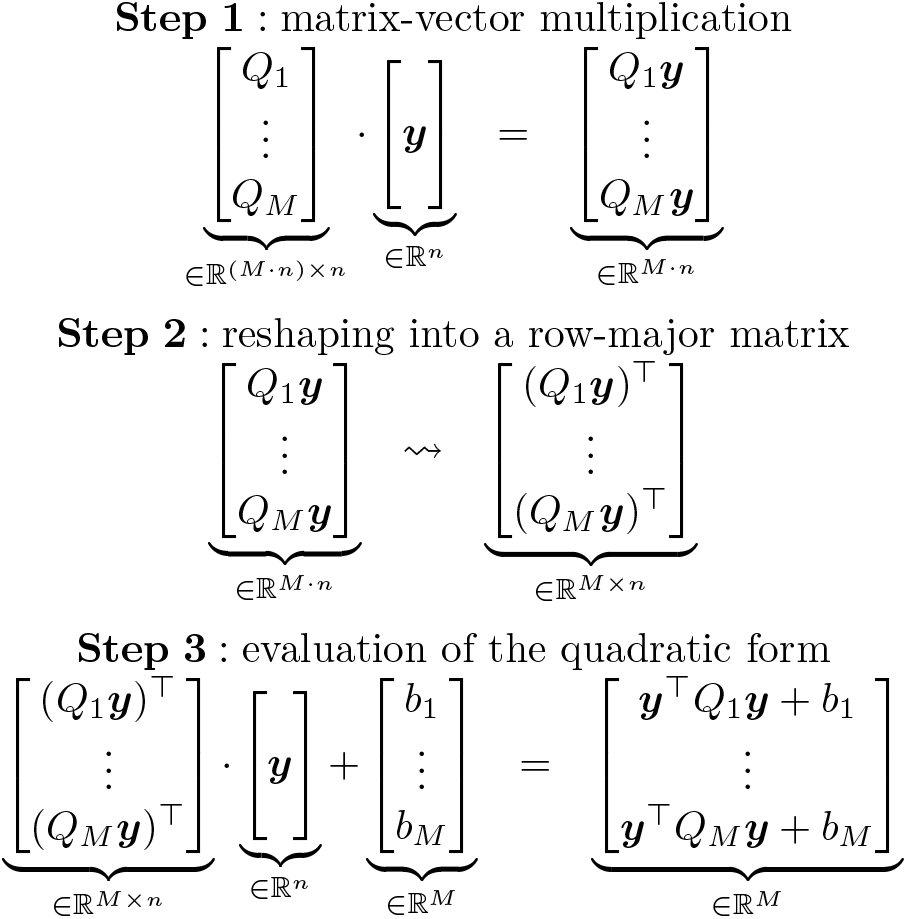
A GPU-accelerated pipeline for the evaluation of quadratic constraints.

## A.5 Accelerated replicate sampling

Figure 3 shows how to accelerate the computation of the constraint verification, by reformulating the *M* quadratic constraints ***y***^T^*Q_m_**y*** + *b_m_* ≥ 0 for *m* = 1, …, *M* in matrix form.

## A.6 Statistical validity

To demonstrate the statistical validity of the kernelPSI approach on GWAS data, we consider the following simulation setup with a genotype matrix *X* ∈ {0, 1, 2} ^*n*×*p*^ of *n* = 100 samples and *p* = 50 SNPs, partitioned in *S* = 10 disjoint and mutually independent subgroups of *p*′ = 5 SNPs. The genotype matrix *X* is sampled as *X* = *X*_1_ + *X*_2_, where the variables in *X*_1_*, X*_2_ ∈ {0, 1} ^*n*×*p*^ are drawn from a Bernoulli distribution with a parameter *p* = 0.4 and with a covariance matrix *V_ij_* = *ρ*^|*i*−*j*|^, *i, j* ∈ {1, · · ·, *p*^t^}. We set the correlation parameter *ρ* to 0.6. To each group corresponds a local identical-by-state kernel *K_i_*.

The outcome *Y* is drawn as *Y* = *θK*_1:3_*U*_1_ + *ϵ*, where *K*_1:3_ = *K*_1_ + *K*_2_ + *K*_3_, *U*_1_ is the eigenvector corresponding to the largest eigenvalue of *K*_1:3_, and *ϵ* is Gaussian noise centered at 0. We vary the effect size of *θ* across the range *θ* ∈ {0.0, 0.01, 0.02, 0.03, 0.05, 0.07, 0.1}, and resample *Y* 1500 times to create 1500 simulations

**Figure S4:**
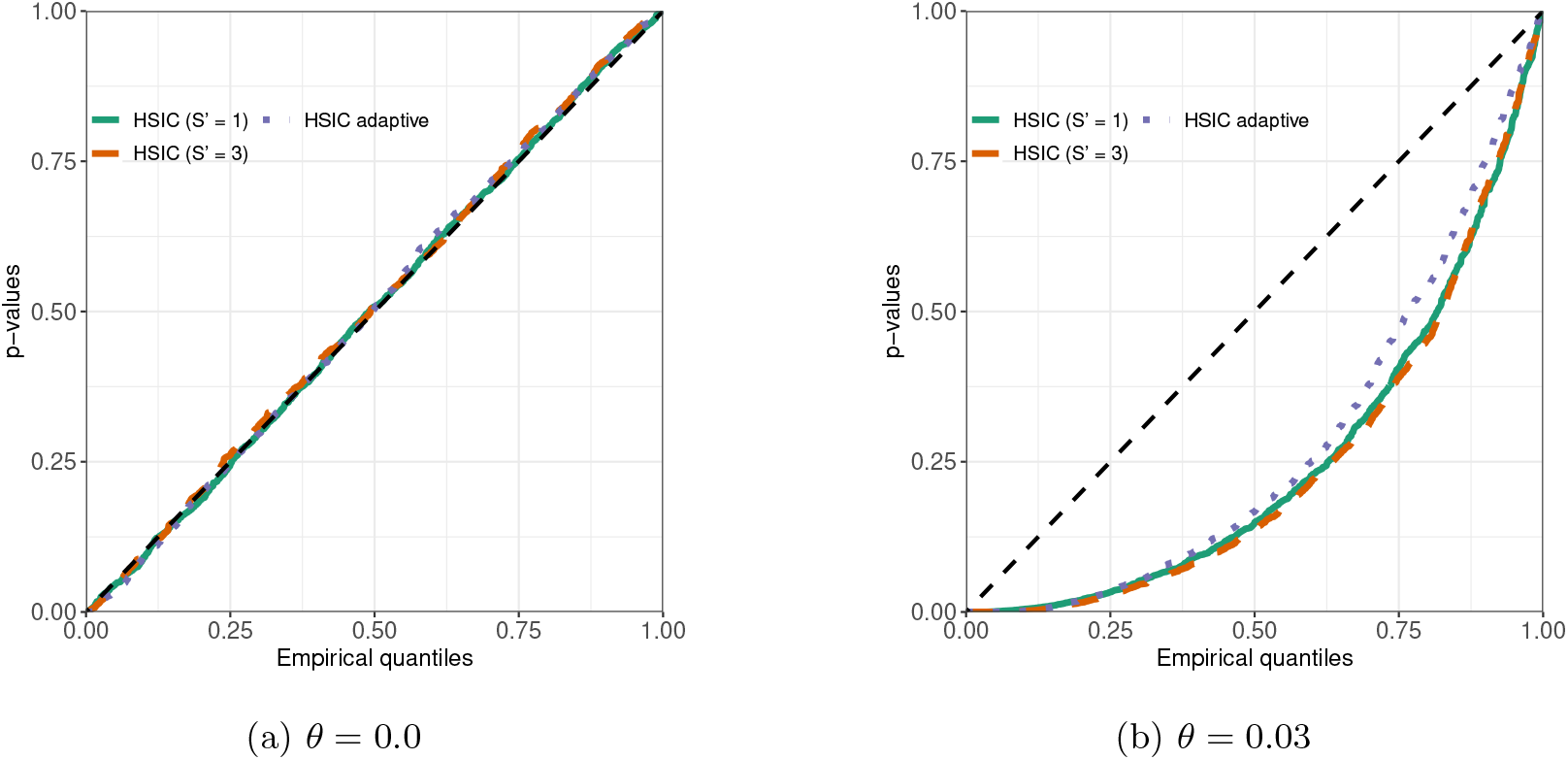
Q-Q plots comparing the empirical kernelPSI p-values distributions under the null hypothesis (*θ* = 0.0) and the alternative hypothesis (*θ* = 0.03) to the uniform distribution.

In Fig 4, we give the results for three different kernel selection strategies: only selecting the most associated kernel (*S*′ = 1), selecting the three most associated kernels (*S*′ = 3), and adaptively determining the number of selected kernels (HSIC adaptive). The latter strategy is also applied for the BMI use-case in our manuscript. Under the null hypothesis *θ* = 0.0, the *p*-values are uniformly distributed for the three kernel selection strategies. This shows that the quadratic kernel association scores in kernelPSI are unbiased. Under the alternative hypothesis *θ* = 0.03 in Figure 4, all kernelPSI variants, particularly adaptive HSIC, have statistical power. This is reflected by low *p*-values and data points located in the bottom-right side of the plot.

**Figure S5:**
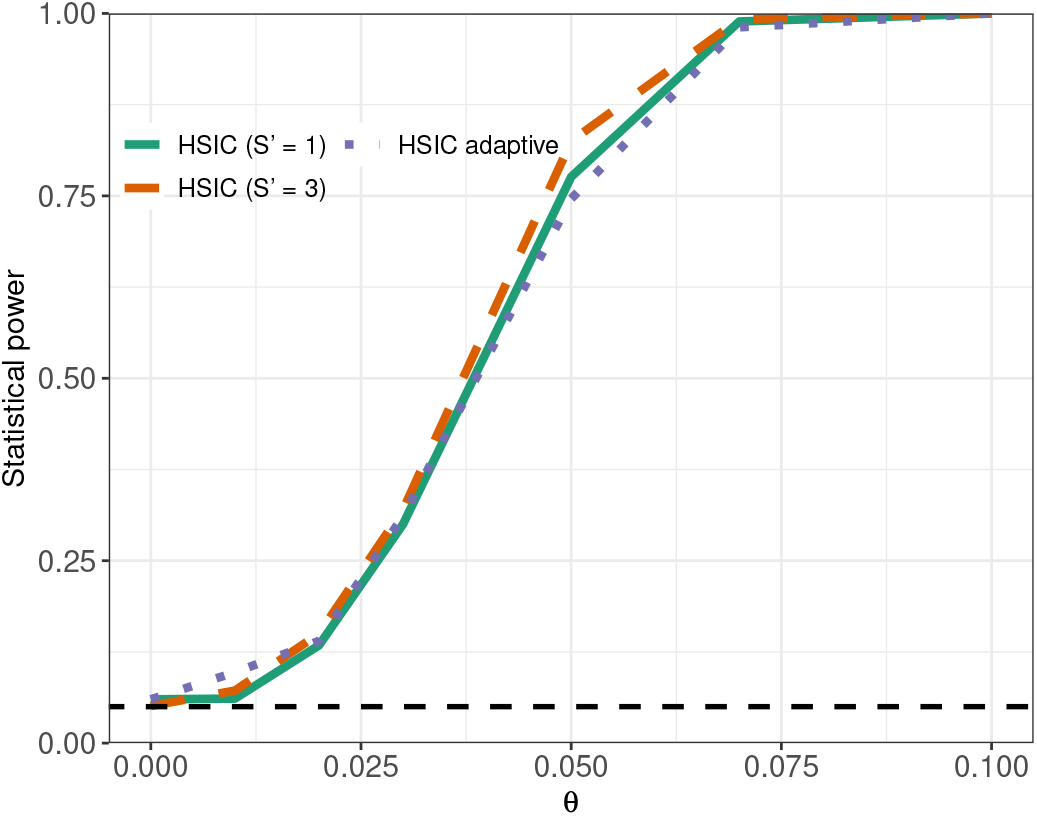
Statistical power of kernelPSI variants, using IBS kernels for simulated genotypic data.

For a significance threshold *α* = 0.05, Figure 5 shows that statistical power increases with the respect to the amplitude of the effect *θ*. Notably, kernelPSI achieves a statistical power of 1 for *θ* = 0.1, and this further confirms the validity of the kernelPSI approach and its capacity in retrieving relevant SNPs and genes.

## A.7 Quality control

The quality control procedure we applied to the UK BioBank dataset is as follows:

- Heterozygosity and missing rates: we discard samples identified by UKBiobank as outliers for both criteria.
- Sex chromosome aneuploidy: only individuals with sex chromosome configurations XX or XY are retained.
- Prior use in phasing: we discard all samples not used in the phasing of autosomal chromosomes.
- Kinship to other participants: we only kept participants with no identified relatives in the dataset.
- Ethnic grouping: we only kept samples identified as “white British” to avoid any potential population structure effects.
- Prior use in principal components analysis (PCA): we discarded all samples not included in the PCA. The analysis is used for population stratification (see next step).
- Homogeneity: additional population structure artifacts are detected used genomic dispersion (GD). We approximated it through the normalized squared distance of the first six principal components (PCs):

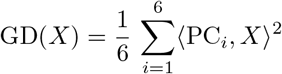

All samples such that GD(*X*) > 3 are considered as outliers and are filtered from the final dataset.

We directly extracted the SNPs of the UK BioBank Axiom array from the imputed genotypes provided by the UK BioBank consortium. As for SNP quality control, we focused on bi-allelic SNPs located on autosomal chromosomes. Moreover, we filtered out the SNPs with a MAF < 0.01, or that are not in Hardy-Weinberg equilibrium (*p* < 1e −10). Out of caution, we also incorporated two additional filters: we shed the SNPs with an internal UK BioBank information score < 0.8 and a missing proportion rate > 1/*n*.

## Footnotes

a More details are provided on their website https://www.nealelab.is/uk-biobank

## Notes

### Competing Interest Statement

The authors have declared no competing interest.

### Summary of Updates

Preprint of an article published in Pacific Symposium on Biocomputing, 2021 World Scientific Publishing Co., Singapore, http://psb.stanford.edu/.

